# KORUS: probabilistic ancestry inference beyond variant calls

**DOI:** 10.64898/2026.11.16.767070

**Authors:** René L Warren, Lauren Coombe, Inanc Birol

## Abstract

Ancestry inference typically relies on variant calling and predefined population labels. We present KORUS, a probabilistic framework that infers ancestry from sequencing-derived k-mer representations constructed using ancestry-informative reference variants and encoded in Bloom filters, without requiring explicit variant calls for query genomes. KORUS separates genomic feature representation from ancestry decoding, enabling supervised classification, fractional ancestry inference, and unsupervised population structure discovery within a unified framework. Reference models were constructed from 643 genomes from the 1000 Genomes Project and the Simons Genome Diversity Project. Evaluation on an independent cohort of 22 whole-genome sequencing samples, with comparisons to ntRoot, SNVstory, and ADMIXTURE, showed recovery of continental-scale population structure, with supervised classification accuracy of 72.7-81.8% and ancestry-fraction concordance of R² = 0.844-0.993 relative to ADMIXTURE and R² = 0.665-0.983 relative to ntRoot. Ancestry estimates were consistent across alternative k-mer representations. Analyses of ancient human and archaic hominin genomes demonstrated that probabilistic projections quantify affinity to the learned feature space rather than necessarily indicating modern continental ancestry. KORUS provides a unified approach to ancestry inference directly from sequencing-derived representations and is freely available under the GPLv3 license at https://github.com/BirolLab/korus.

## Introduction

Human genomic variation is structured by population history, migration, isolation, and admixture. Characterizing this structure has become a central component of modern genomic research, where ancestry information is routinely used to study demographic history, interpret genetic variation, control population stratification, and contextualize biomedical findings^1,2^. As population-scale sequencing projects and biobank resources have expanded, ancestry inference has evolved from a specialized population genetics application into a routine analytical component of large-scale genomic studies^3–5^.

Most contemporary ancestry inference methods operate within a variant-centric paradigm. Sequencing reads are aligned to a reference genome, genetic variants are identified, and ancestry is subsequently inferred from genotypes, haplotypes, or population-specific allele frequencies. This paradigm underlies widely used approaches including STRUCTURE^6^, ADMIXTURE^7^, EIGENSTRAT^8^, RFMix^9^, FLARE^10^, and numerous related local and global ancestry inference frameworks, including ntRoot^11^. Although these methods differ substantially in their statistical models and computational approaches, they share a common dependence on explicit variant representation. The success of variant-based ancestry inference has been enabled by reference resources such as the International HapMap Project^12^, the 1000 Genomes Project^1^, and the Simons Genome Diversity Project^2^, which collectively established large catalogues of human genetic variation across diverse populations. However, the resulting ancestry labels and population assignments represent abstractions of complex demographic histories rather than discrete biological categories. Recent recommendations from the National Academies have emphasized the limitations of treating ancestry as a fixed categorical attribute and have encouraged more nuanced representations of human genetic diversity^13^. These considerations motivate a broader question: to what extent is ancestry inference intrinsically dependent upon explicit sample variant identification? From the perspective of genomic data representation, the process of variant calling represents a substantial dimensionality reduction of the original sequencing data. Billions of observed nucleotides are transformed into a comparatively small collection of variant states that are subsequently used for ancestry estimation. While highly effective, this workflow assumes that explicit sample variant calls provide the optimal representation for ancestry inference.

Recent advances in alignment-free genomics approaches suggest alternative representations may also be informative. K-mer–based methods have demonstrated utility across genome assembly, sequence comparison, metagenomics, and variant detection, owing to their scalability and ability to operate directly on sequencing reads^14–16^. Previous work demonstrated that population structure can be detected directly from k-mer content without explicit genotype calling^17^, indicating that ancestry-related information is embedded within sequence composition itself. Building on these observations, ntRoot introduced a reference-guided, alignment-free framework for ancestry inference based on Bloom filter interrogation of ancestry-informative variants^11^. Rather than relying on sequence alignment, ntRoot identifies known variants directly from sequencing-derived k-mer representations, enabling accurate recovery of both global and local ancestry with high concordance to conventional methods. Although ntRoot eliminated the requirement for alignment and conventional genotyping, it remains fundamentally variant-centric because ancestry estimation is still derived from explicit single nucleotide variant identification.

Here, we present KORUS (K-mer Orthogonal Representation for Unified Stratification), a framework that extends this alignment-free paradigm by performing ancestry inference directly from genome-wide probabilistic feature representations rather than explicit sample variant calls. Instead of first identifying variants and subsequently estimating ancestry, KORUS treats ancestry inference as a probabilistic decoding problem over genomic feature representations. Population-informative sequence features are aggregated into a genome-wide feature representation derived directly from sequencing data, and ancestry is inferred from the structure of that representation rather than from discrete variant calls. This formulation allows ancestry information to be modeled using either categorical population labels, continuous ancestry fractions, or entirely unsupervised representations, thereby reducing dependence on predefined ancestry classes. KORUS reference models were constructed from whole-genome sequencing data representing globally distributed populations from the 1000 Genomes Project and Simons Genome Diversity Project. We evaluated supervised, fraction-supervised, and fully unsupervised ancestry inference using an independent blind-test cohort and compared performance with ntRoot^11^, SNVstory^18^, and ADMIXTURE^7^. We further examined the behaviour of the framework on ancient human and archaic hominin genomes to explore probabilistic projections beyond the modern reference distribution.

## Results

### Overview of the KORUS ancestry inference framework

KORUS performs ancestry inference from sequencing-derived k-mer signals using a two-stage computational framework that transforms allele-specific genomic signals into probabilistic ancestry assignments (Fig. 1). First, ancestry-informative k-mers, derived from ancestry-associated allele frequency differences in the GRCh38-integrated 1000 Genomes reference panel^11,19,20^ (Supplementary Table 1), are queried directly against Bloom filters generated from sample sequencing reads. These k-mers capture sequence contexts containing either the reference or alternative allele at informative variant sites, producing normalized allele-specific signals that are aggregated across fixed genomic windows to generate a sample-by-window feature matrix. The resulting genomic representation is standardized and projected into an orthogonal principal component (PCA) feature space prior to ancestry modelling.

**Figure 1.**
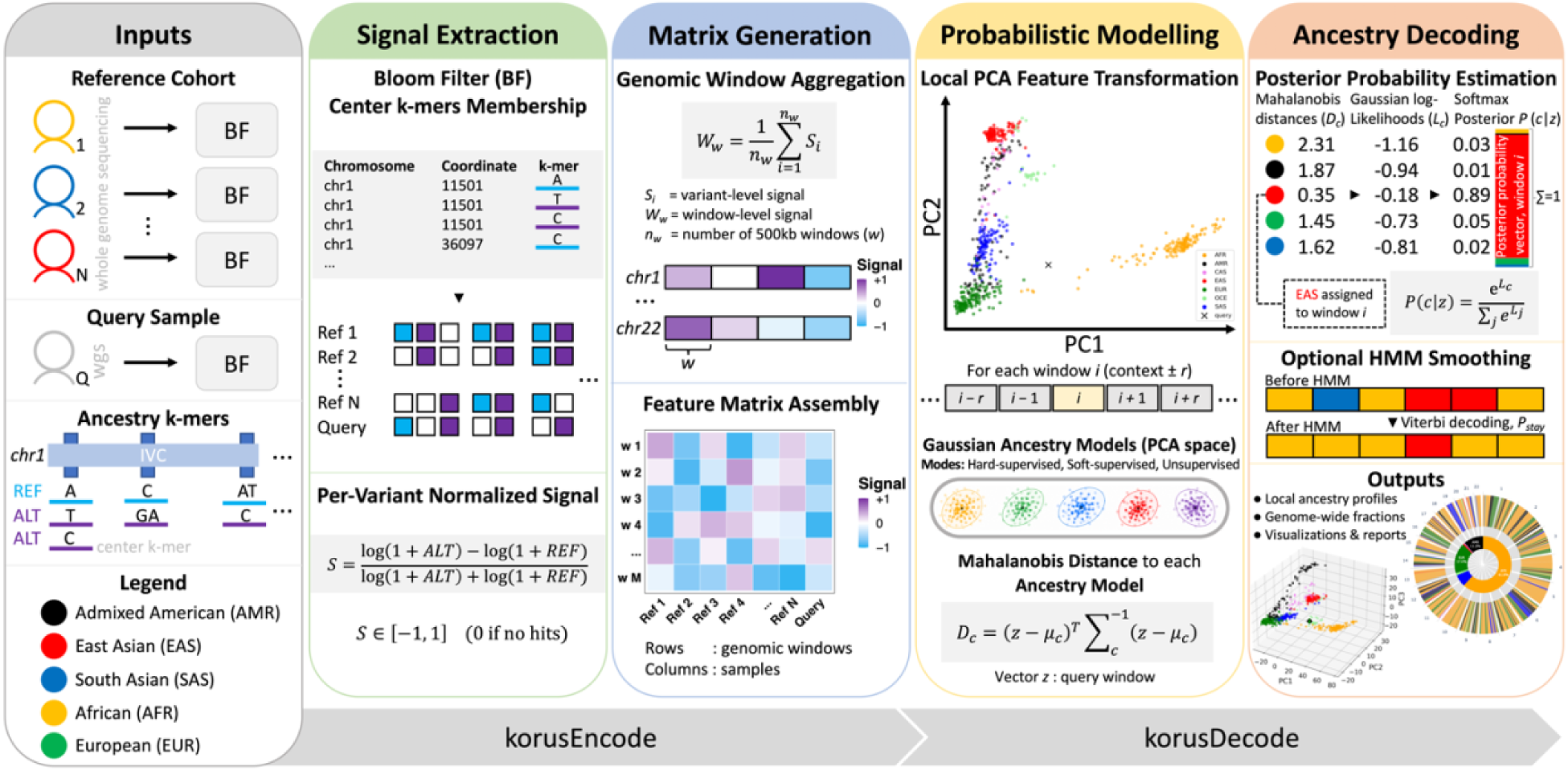
Overview of the KORUS (K-mer Orthogonal Representation for Unified Stratification) workflow. **Inputs**. Whole-genome sequencing (WGS) reads from reference and query genomes are converted into Bloom filters (BF) and interrogated using ancestry-informative REF and ALT allele k-mer catalogues derived from the integrated variant catalogue (IVC). **Signal Extraction**: sequencing-derived BFs from reference and query genomes are interrogated using ancestry-informative REF and ALT allele k-mers to quantify reference- and alternate-allele sequence support. Per-locus normalized REF–ALT signals are computed and aggregated within fixed genomic windows. **Matrix Generation**: window-level ancestry signals are averaged to generate genome-wide ancestry signal profiles, which are compiled into a genomic feature matrix in which rows correspond to genomic windows and columns correspond to genome samples. **Probabilistic Modelling**: the feature matrix is standardized and projected into a local principal component (PCA) feature space to reduce feature correlation and improve covariance estimation. For each genomic window, ancestry likelihoods are estimated using Mahalanobis distance to Gaussian models estimated from either predefined ancestry labels (hard-supervised), fractional ancestry supervision (soft-supervised), or unsupervised K-means-derived pseudo-labels. **Ancestry Decoding**: Mahalanobis distances are converted into posterior ancestry probabilities by softmax normalization and may optionally be refined using Hidden Markov Model (HMM) smoothing with Viterbi decoding to enforce local genomic continuity. Final outputs include genome-wide ancestry fractions, per-window local ancestry predictions with posterior probabilities, interactive PCA visualizations, and local ancestry reports.

Window-level ancestry is then inferred using Mahalanobis distance to ancestry-specific Gaussian models estimated from either discrete ancestry labels, fractional ancestry proportions, or unsupervised K-means-derived pseudo-labels. Distances are converted to posterior ancestry probabilities through softmax normalization, and optional HMM smoothing enforces genomic continuity of local ancestry predictions. The framework simultaneously produces local ancestry assignments, genome-wide ancestry fractions, probabilistic ancestry estimates, and interactive visualizations of both local and global ancestry relationships.

### KORUS recapitulates global population structure across multiple ancestry inference modes

To evaluate the flexibility of the probabilistic framework, KORUS was applied to a globally diverse reference panel of 643 genomes representing seven broad ancestry groups (see Methods; Supplementary Table 2), using hard-supervised, unsupervised, and soft-supervised ancestry inference. Despite their differing supervision strategies, all three approaches recovered the major population structure present in the reference panel (Fig. 2).

**Figure 2.**
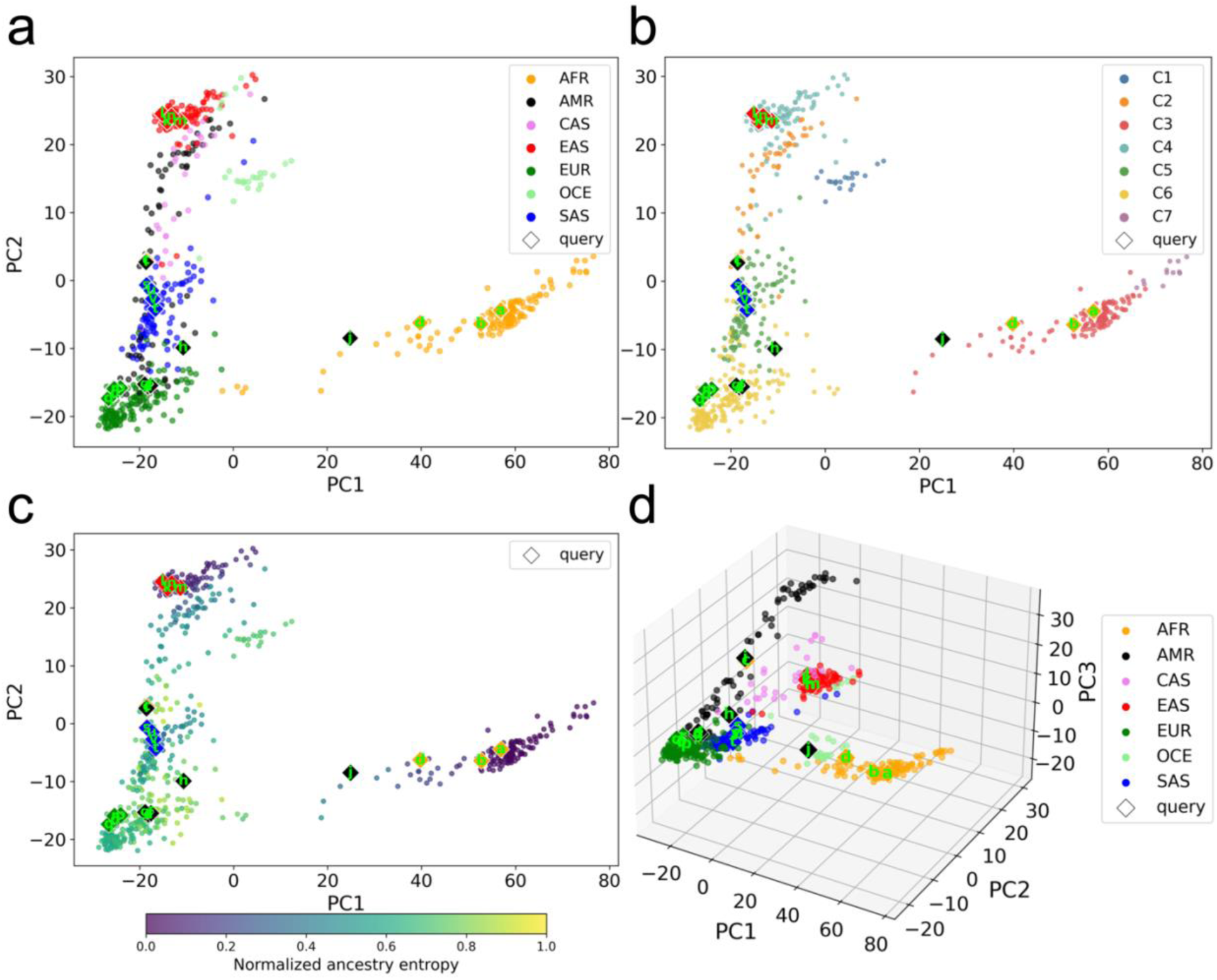
Principal component representations generated by KORUS under three ancestry inference modes. **(a)** Hard-supervised decoding using discrete ancestry labels produces well-separated continental ancestry clusters, with query genomes shown as diamonds colour-coded by their 1kGP superpopulation annotations and projected into the learned ancestry space. **(b)** Unsupervised decoding partitions the reference panel into seven K-means clusters without prior ancestry labels. Cluster assignments closely recapitulate major continental population structure while permitting finer subdivision of genetically distinct groups. **(c)** Soft-supervised decoding models ancestry using fractional reference memberships, yielding a continuous ancestry manifold in which admixed reference genomes occupy intermediate positions between major ancestry groups. Reference genomes are coloured according to normalized ancestry entropy, with darker colours indicating lower entropy (more homogeneous ancestry) and lighter colours indicating higher entropy (greater ancestry admixture). **(d)** Three-dimensional PCA visualization showing the reference cohort alongside projected blind-test query genomes (diamond symbols), colour-coded by their original 1kGP-assigned superpopulation labels. The additional principal component improves visualization of population separation while providing population context for projected query genomes.

Under hard supervision, discrete ancestry labels produced well-separated continental clusters within PCA space, with African, East Asian, South Asian, Oceanian, European, and Admixed American populations forming distinct groups while positioning each query genome within the appropriate ancestry context (Fig. 2a; Supplementary Table 3). This representation reflects the covariance structure learned from categorical ancestry labels and provides the reference framework used for supervised local ancestry inference.

KORUS also supported completely unsupervised ancestry inference through K-means clustering of reference genomes in PCA space (Fig. 2b). Without providing ancestry labels or ancestry fractions, unsupervised clustering recovered comparable major population structure observed under supervised analyses while allowing genetically related populations to organize naturally according to genomic similarity. Importantly, the inferred cluster assignments were used only to initialize probabilistic ancestry modelling, demonstrating that KORUS can operate independently of predefined ancestry categories.

Replacing discrete labels with fractional ancestry supervision generated a continuous ancestry representation in which admixed reference genomes occupied intermediate positions between major ancestry groups (Fig. 2c). Rather than assigning each reference genome to a single population, fractional supervision estimated ancestry-specific Gaussian models using weighted ancestry contributions from each individual, producing a smoother ancestry manifold that better reflects the continuous nature of human genetic diversity. The accompanying entropy visualization illustrates the expected increase in ancestry uncertainty among admixed individuals relative to genetically homogeneous populations.

To complement these ancestry-specific representations, KORUS also generates three-dimensional PCA visualizations of the reference cohort together with projected query genomes (Example for hard-supervision in Fig. 2d). The additional dimension improves visual separation of major ancestry groups while providing the population-context view incorporated into the interactive HTML report described below. Blind-test samples project alongside their corresponding reference populations, illustrating the consistency of the learned genomic representation across independent datasets.

Together, these analyses demonstrate that the underlying genomic feature representation is sufficiently informative to recover global population structure under three distinct probabilistic inference paradigms while providing a unified framework for discrete, continuous, and entirely label-free ancestry modelling.

### Ancestry classification accuracy

To assess the accuracy of KORUS ancestry classifications, we compared KORUS ancestry fraction estimates with those from ntRoot, ADMIXTURE, and SNVstory using 22 human samples (Supplementary Tables S4-S9). KORUS achieved accuracies of 72.7% and 81.8% using hard- and soft-supervised inferences, respectively. In comparison, ntRoot, ADMIXTURE and SNVstory yielded accuracies of 90.9%, 90.9% and 100.0%, respectively, demonstrating how the KORUS framework can generate similarly accurate results without using variant calling. Even when KORUS was run in unsupervised mode, thus not using any reference labels, it still achieved an accuracy of 68.2%. We note that SNVstory uses a model built from 1kGP samples, therefore the query samples in the blind set may be in their training set and thus inflate the accuracy statistics for this tool. We also computed a normalized multiclass Brier score for each tool, which takes the computed ancestry fractions into account (Table 1). KORUS yielded Brier scores of 0.14-0.24, while ntRoot and ADMIXTURE classifications were more concordant with the ground truth labels, as indicated by lower Brier scores (0.13 and 0.10, respectively). Because SNVstory does not output estimated ancestral fractions, the multiclass Brier score could not be computed for this tool.

**Table 1.**
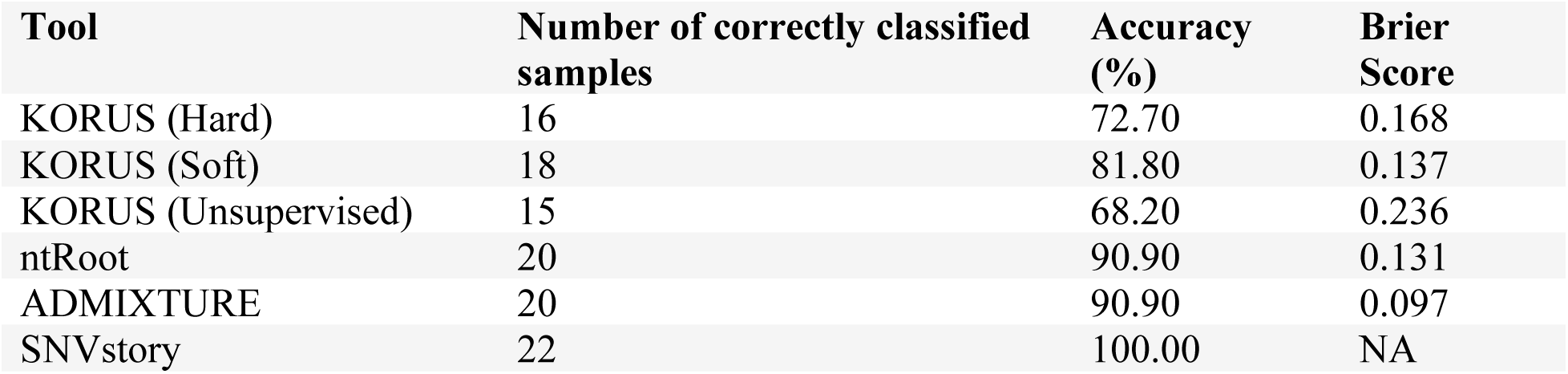
Accuracy and normalized multiclass Brier scores from ancestry predictions using KORUS, ntRoot, ADMIXTURE and SNVstory. Global ancestry inference was performed using whole genome sequencing reads from 22 human samples. For KORUS (k=55), ntRoot and ADMIXTURE, the global ancestry label was assigned to the super-population with the largest ancestry fraction, while for SNVstory, the label with the highest probability was chosen. The classification was considered correct if it matched the 1kGP label, and the accuracy was calculated as the percentage of correctly classified samples.

Notably, all tools except for SNVstory misclassified samples HG01243 and NA20314, based on the 1kGP ground truth (Supplementary Table 10). A previous study by Zimin *et al.* (2022) found that while the 1kGP consortium labelled HG01243 as Admixed American, the genomic data indicated that the individual had predominantly African ancestry^21^. Similarly, NA20314 is labelled as African by 1kGP, but several studies have found that the sample has limited African ancestry, and groups better with Admixed American samples^22–24^. If the ground truth labels for these samples are adjusted based on these past studies, the accuracy of the KORUS predictions in hard and soft supervised modes increases to 81.8-90.9% (Supplementary Table 11). After this adjustment, the accuracies for ntRoot and ADMIXTURE both increase to 100%, while the SNVstory accuracy drops to 90.9%.

### Quantitative comparison of ancestry fraction estimates across inference modes and reference methods

To quantitatively assess the robustness of KORUS ancestry estimates, we compared genome-wide ancestry fractions obtained under hard- and soft-supervised inference, evaluated reproducibility across two independent k-mer sizes (*k* = 45 and *k* = 55), and benchmarked KORUS against the previously developed alignment-free ancestry framework ntRoot as well as ADMIXTURE (Fig. 3, Supplementary Table 12). Comparisons between hard- and soft-supervised KORUS models demonstrated strong overall concordance, indicating that both supervision strategies recovered highly consistent ancestry profiles despite their different modelling assumptions (R^2^ values: 0.908-0.996) (Fig. 3a). Minor systematic differences were nevertheless apparent. Hard-supervised inference tended to assign slightly larger European ancestry fractions, whereas soft-supervised inference more frequently allocated modest increases to Admixed American ancestry components. These differences were primarily observed among admixed individuals and likely reflect the ability of fractional supervision to model continuous ancestry structure without constraining reference genomes to single ancestry labels.

**Figure 3.**
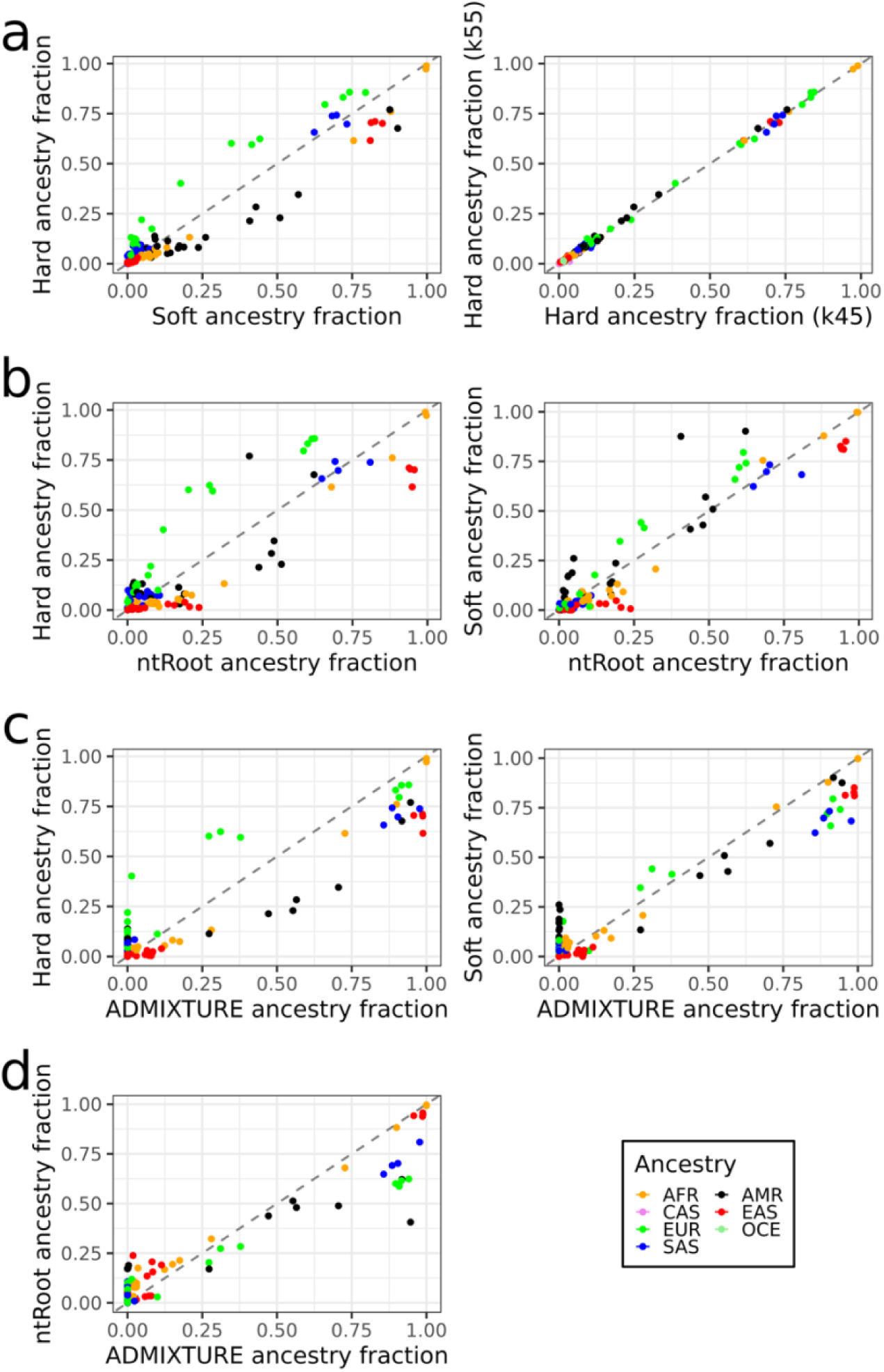
Quantitative validation of KORUS ancestry estimates including concordance of genome-wide ancestry fractions estimated by KORUS relative to ntRoot and ADMIXTURE. **(a)** Comparison of hard-supervised and soft-supervised KORUS ancestry fractions demonstrates high overall agreement, with modest systematic differences attributable to the supervision strategy. Hard-supervised inference slightly increases European ancestry estimates, whereas fractional supervision tends to assign modestly higher Admixed American (AMR) ancestry proportions. **(b)** Comparison of KORUS in hard-supervised mode with ntRoot using k-mer sizes of 45 and 55 demonstrates highly consistent ancestry estimates across both parameterizations. Dominant ancestry assignments remained unchanged, with only minor quantitative differences in secondary ancestry fractions. **(c)** Comparison of KORUS in hard- and soft-supervised modes with ADMIXTURE and **(d)** comparison of ntRoot and ADMIXTURE similarly shows strong concordance across ancestry fractions. Across all methods, differences attributable to k-mer length are substantially smaller than method-specific differences between inference frameworks, indicating that ancestry estimation is robust to k-mer size while reflecting intrinsic characteristics of each probabilistic model.

KORUS also exhibited high robustness to k-mer size. Ancestry fractions inferred using models constructed from *k* = 45 and *k* = 55 ancestry-informative k-mers were very similar for both hard- and soft-supervised analyses (Fig. 3a, Supplementary Table 12), with R^2^ values ranging from 0.708-1.000. Across all samples, only minor quantitative differences in ancestry fractions were observed, typically involving small adjustments to secondary ancestry components, while dominant ancestry assignments and the overall population structure remained unchanged. These findings indicate that ancestry inference is largely insensitive to moderate changes in k-mer length, suggesting that the probabilistic decoding framework is driven primarily by the underlying genomic ancestry signal rather than the precise k-mer representation. Comparison with ntRoot demonstrated similarly high overall concordance (R^2^ values 0.665-0.976 and 0.825-0.983 for hard- and soft-inference, respectively) (Fig. 3b; Supplementary Table 12). For individuals with a strongly predominant ancestry component, both methods consistently identified the same predominant ancestry irrespective of k-mer size (Supplementary Fig. 1). Individuals with more distributed ancestry profiles showed greater variation in inferred ancestral fractions across k-mer sizes for ntRoot than for KORUS. This was particularly evident for the European and Admixed American samples from their deviations from the *y=x* line and their corresponding concordance correlation coefficient (CCC) values (ntRoot: 0.961 and 0.991; KORUS: 0.999 and 0.998 for EUR and AMR, respectively) (Fig. 3a, Supplementary Fig. 1, Supplementary Table 12). Importantly, these systematic differences were substantially larger than those introduced by changing k-mer size, indicating that methodological differences between the probabilistic inference frameworks dominate over differences arising from the underlying sequence representation. Benchmarking against ADMIXTURE yielded similarly strong correlations (R^2^ values 0.844-0.989 and 0.894-0.993 for hard- and soft-inference, respectively), further supporting the accuracy of KORUS ancestry estimates across diverse population backgrounds (Fig. 3c). Although individual ancestry fractions differed modestly among methods, particularly for highly admixed genomes, the overall ancestry structure was consistently recovered. Collectively, these analyses demonstrate that KORUS produces stable and reproducible ancestry estimates across multiple k-mer representations while achieving strong agreement with established ancestry inference approaches.

### Unsupervised clustering recovers population structure

To evaluate whether KORUS could recover ancestry structure without prior population labels, reference genomes were clustered using K-means in PCA space and the resulting cluster assignments were subsequently used as pseudo-labels for probabilistic ancestry modelling (*k* = 55; Fig. 4). Unsupervised clustering consistently recovered the major continental-scale population structure of the reference panel. Three clusters exhibited complete purity with respect to the known reference ancestry labels, corresponding to Oceanian and two African subclusters, while the remaining clusters were strongly enriched for South Asian (92.2% purity), Admixed American (85.4%), East Asian (84.7%), and European (75.4%) ancestries (Fig. 4).

**Figure 4.**
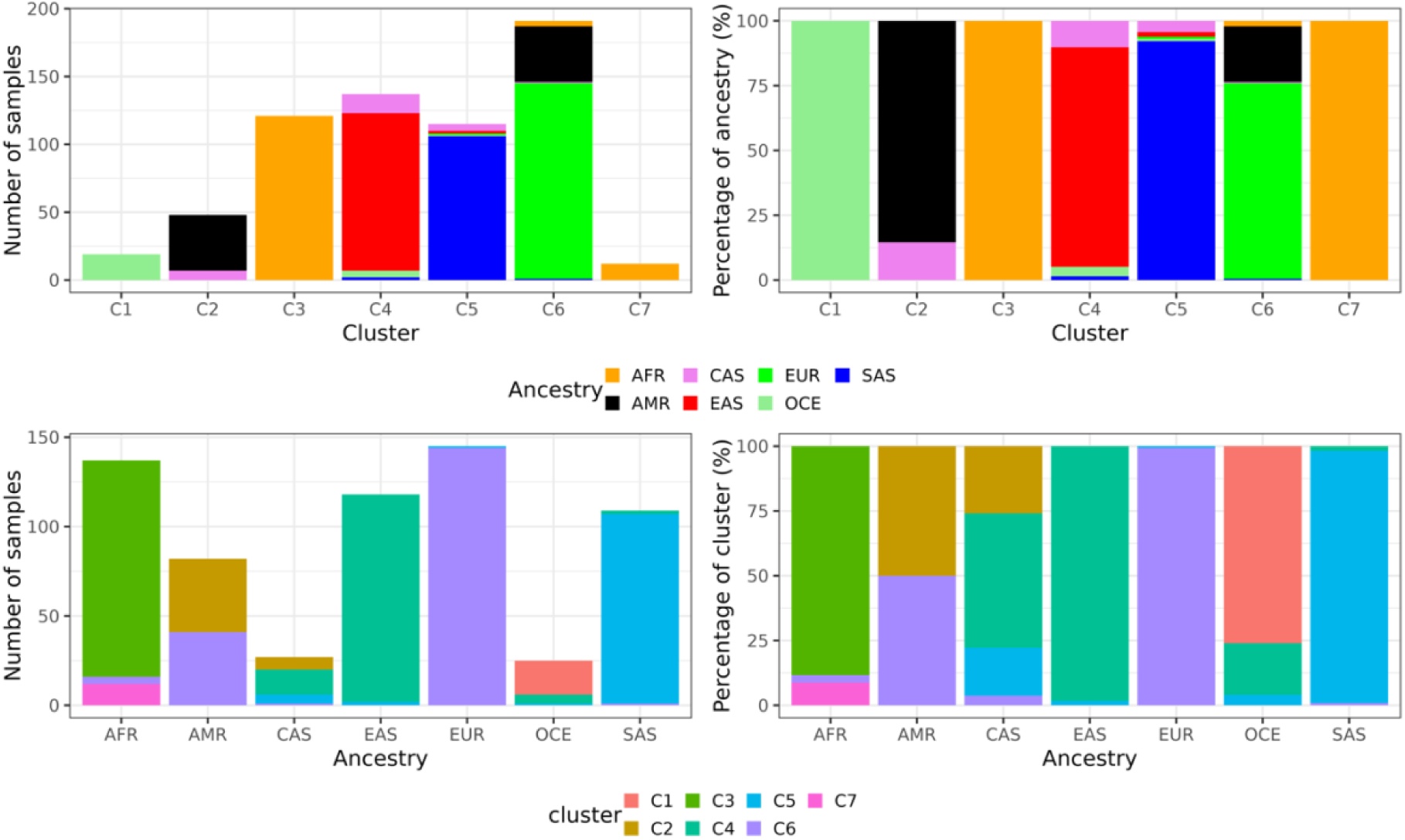
Unsupervised recovery of population structure by KORUS. Characterization of unsupervised clustering of the reference panel using seven K-means clusters. Cluster composition is summarized for independent analyses using a k-mer length of 55. Stacked bar plots illustrate the distribution of known reference ancestries within each inferred cluster and the reciprocal distribution of ancestry groups across clusters. Several clusters correspond almost exclusively to single continental populations, including African and Oceanian clusters exhibiting 100% purity, whereas the European-predominant cluster contains the greatest ancestral heterogeneity, reflecting the genetic proximity of European, Admixed American, and Central Asian reference populations. Admixed American and Central Asian reference genomes are distributed across multiple inferred clusters, consistent with their greater genetic diversity and admixed ancestry.

Further inspection of cluster composition revealed biologically interpretable patterns (Fig. 4). All European reference genomes localized to a single dominant cluster, although this cluster also contained a subset of Admixed American samples together with a small number of African, Central Asian, and South Asian individuals. Similarly, Admixed American reference genomes were partitioned between a predominantly Admixed American cluster and the European cluster, consistent with the documented European ancestry contributions among Admixed American populations represented in the 1000 Genomes Project^1,25^. Central Asian genomes likewise did not form an independent cluster but were distributed primarily among the South Asian, East Asian, and Admixed American enriched clusters, reflecting their intermediate genetic relationships to neighboring populations rather than constituting a distinct continental-scale lineage.

Perhaps the most notable observation was the partitioning of African reference genomes into two completely pure clusters. Rather than representing clustering instability, this result likely reflects genuine population substructure within the African reference panel. African populations harbor the greatest levels of human genetic diversity^26^, and the separation into two high-purity clusters suggests that KORUS is sufficiently sensitive to capture finer-scale genomic structure while remaining entirely agnostic to predefined ancestry labels. Together, these findings demonstrate that the unsupervised implementation recovers biologically meaningful population organization directly from genomic feature space, supporting its application to reference collections where ancestry labels are incomplete, uncertain, or unavailable.

To assess the impact of k-mer size on the unsupervised clustering, the analysis was also performed using *k* = 45 (Supplementary Fig. 2). Nearly identical clustering patterns were obtained for both k-mer sizes, and K-means clustering converged to identical solutions across independent centroid initializations (n_init = 10), demonstrating a high level of reproducibility of the unsupervised framework. The correspondence between clusters obtained using *k* = 45 and *k* = 55 further illustrated the stability of the learned ancestry structure. Most clusters mapped directly between the two models, including the principal African, European, Oceanian, and secondary African clusters.

### Interactive visualization integrates global and local ancestry inference

In addition to genome-wide ancestry estimates, KORUS generates an interactive HTML report that combines chromosome-scale local ancestry inference with global population structure visualization into a unified analysis environment (Fig. 5). The report integrates the circular local ancestry plot produced by the ntRoot visualization framework with an interactive three-dimensional PCA representation of the reference cohort and query genome generated directly by KORUS. Linked interactions between the two views allow users to simultaneously examine ancestry assignments along the genome while visualizing the position of the query genome relative to the reference population.

**Figure 5.**
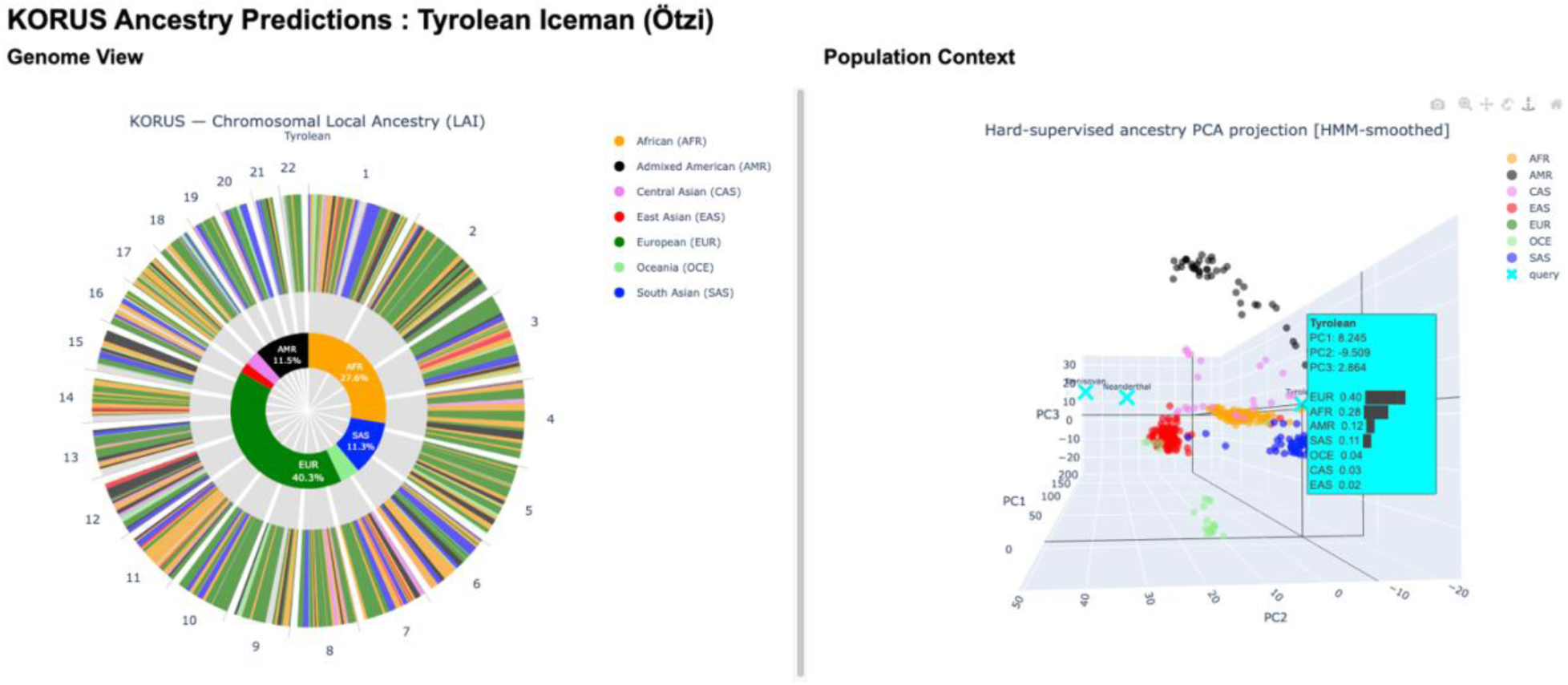
Example interactive HTML report generated by the end-to-end KORUS pipeline for the Tyrolean Iceman (Ötzi). The report integrates chromosome-scale local ancestry inference (left) with an interactive three-dimensional PCA representation of the reference population and query genome (right). Local ancestry is visualized across all chromosomes using circular genome plots, while the PCA display provides population context for the inferred ancestry profile. Interactive highlighting links the global population projection with corresponding ancestry segments in the genome view, facilitating simultaneous exploration of genome-wide ancestry composition and local ancestry transitions. The report is generated automatically for each analyzed sample and is designed to support both routine ancestry analyses and exploratory investigation of complex genomes. The interactive report is available at: https://www.birollab.ca/viz/fig5_TyrHardKor45-ancient.report.html

To illustrate the utility of the integrated report, we analyzed the genome of the Tyrolean Iceman (Ötzi), a ∼5,300-year-old Copper Age individual from the eastern Italian Alps (Fig. 5; Supplementary Fig. 3). The inferred ancestry profile was dominated by European ancestry, consistent with previous genomic analyses showing that Ötzi derives from ancestry components related to ancient and present-day European populations^27–29^, while also identifying lower-probability ancestry components that reflect the probabilistic nature of ancestry estimation within the learned ancestry manifold. The interactive PCA view simultaneously places the ancient query genome within the modern reference population manifold, enabling local ancestry patterns to be interpreted within their broader genomic context. A three-dimensional PCA including archaic hominin genomes is provided in Supplementary Fig. 3.

By integrating quantitative ancestry estimates, local ancestry visualization, and interactive exploration of population relationships within a single report, KORUS provides an intuitive interface for interpreting both local and global ancestry simultaneously, facilitating routine ancestry analysis as well as exploratory investigation of complex genomes.

## Discussion

KORUS provides a unified framework for ancestry inference from k-mer-derived genomic representations, separating feature construction from probabilistic inference and population supervision. This modular architecture supports discrete-label, fractional-weight, and unsupervised modelling within a common framework, enabling comparison of alternative ancestry inference strategies using a shared genomic representation.

In many ancestry inference frameworks, PCA is primarily used for visualization or exploratory analyses^30^. In KORUS, however, PCA serves a fundamentally different purpose. By transforming correlated genomic window signals into an orthogonal latent representation, the covariance structure becomes substantially more compatible with Mahalanobis distance estimation. The principal components therefore define the statistical space in which ancestry likelihoods are estimated rather than simply providing a low-dimensional visualization of population structure. The name KORUS (K-mer Orthogonal Representation for Unified Stratification) reflects this central role of orthogonal feature representations within the probabilistic decoding framework. An important implication of these findings is that the genomic representation and the statistical decoder should be viewed as distinct components of ancestry inference. Across two independent k-mer sizes, ancestry estimates remained highly stable despite substantial changes in the underlying sequence representation, whereas systematic differences persisted between KORUS and ntRoot irrespective of k. These observations suggest that, once a sufficiently informative genomic representation has been established, methodological differences arise primarily from the probabilistic model used to interpret that representation rather than from the representation itself. This separation creates opportunities for future methodological development in which alternative probabilistic models can be evaluated independently of feature generation.

Most ancestry inference methods are intrinsically tied to predefined population labels or fixed admixture models^7,9^. KORUS instead separates supervision from representation, allowing ancestry information to be incorporated as discrete labels, continuous ancestry fractions, or omitted entirely. These supervision strategies differ only in how probability distributions are estimated; the underlying genomic representation and probabilistic decoding framework remain unchanged. Consequently, the same computational architecture naturally supports conventional supervised classification, continuous ancestry modelling, and unsupervised discovery of population structure. The high-purity clusters recovered in unsupervised mode further demonstrate that much of the large-scale structure of human genetic diversity is recoverable directly from the learned genomic representation, even in the absence of predefined ancestry labels.

The projection of archaic hominin genomes further illustrates an important property shared by all supervised ancestry inference methods. Because Denisovan and Neanderthal genomes lie well outside the distribution represented by present-day human reference panels, the resulting posterior probabilities quantify affinity to the learned modern genomic manifold rather than biologically meaningful continental ancestry. Similar behavior has been reported for model-based ancestry inference methods, where ancient genomes projected into modern reference panels are represented as combinations of modern ancestry components rather than as distinct ancestral lineages^31^. This behavior highlights the importance of distinguishing interpolation within the training distribution from extrapolation beyond it. Like all reference-based ancestry inference methods, KORUS remains dependent on the diversity and representativeness of the reference panel used for model construction. Population structure that is absent or underrepresented within the reference data cannot be recovered de novo by supervised inference, and the resolution of ancestry estimates is therefore ultimately constrained by the granularity of the available reference populations. Although the present study focuses on continental-scale ancestry using 1000 Genomes-derived reference data^1^, the framework itself is agnostic to the taxonomic resolution of the training panel and can readily accommodate finer-scale regional or population-specific reference datasets as these continue to expand.

More broadly, KORUS suggests that ancestry inference need not ultimately be framed as prediction of predefined population labels. Human genetic diversity is inherently continuous, shaped by migration, admixture, demographic history, and evolutionary divergence^32,33^. Population labels remain valuable for interpretation, benchmarking, and communication, but they are abstractions imposed upon an underlying continuum of genetic relationships^31^. By supporting hard supervision, fractional supervision, and completely unsupervised decoding within a common probabilistic framework, KORUS demonstrates that ancestry can instead be represented as position within a learned genomic manifold whose interpretation is not intrinsically tied to discrete categories. As increasingly diverse global reference resources become available, this separation between representation and supervision may permit ancestry models to evolve from classifiers of predefined populations toward quantitative reconstructions of continuous genomic relationships. In this sense, KORUS is not simply an alternative ancestry classifier. Rather, it provides a general computational framework in which genomic feature representations, probabilistic inference, and population supervision become modular components that can evolve independently as sequencing technologies and reference resources continue to expand. We believe this architecture represents an important step toward ancestry inference beyond labels, consistent with recent efforts to move beyond simplistic population descriptors toward representations that better capture the continuous nature of human genetic diversity^13^.

With KORUS, we demonstrate that ancestry inference can be performed through probabilistic decoding of k-mer-derived genomic representations without explicit alignment-based variant calling. By separating genomic representation from probabilistic inference and population supervision, KORUS accommodates discrete labels, fractional ancestry models, and unsupervised population discovery within a unified framework. As genomic reference resources expand in size and diversity, this modular architecture provides a foundation for investigating ancestry beyond predefined population categories.

## Methods

### Overview of the KORUS framework

KORUS (K-mer Orthogonal Representation for Unified Stratification) is a two-stage framework for ancestry inference from whole-genome sequencing data. Rather than performing read alignment and variant calling, KORUS operates directly on sequencing-derived k-mer representations encoded in Bloom filters. The framework consists of an encoding stage that converts ancestry-informative variant k-mers into a genomic signal matrix and a decoding stage that transforms these signals into probabilistic ancestry assignments across the genome.

The central design principle of KORUS is that ancestry information can be represented as a continuous genomic signal derived from differential support for reference and alternate allele sequence k-mers generated from ancestry-informative loci. These signals are aggregated across genomic windows and interpreted in a covariance-aware probabilistic models in a PCA-derived feature space using probabilistic classification and optional Hidden Markov Model (HMM) smoothing. The workflow consists of five main stages: (i) Bloom filter construction, (ii) allele-specific signal extraction, (iii) genomic window aggregation, (iv) ancestry feature-space construction, and (v) probabilistic ancestry decoding.

The same genomic feature representation supports three complementary probabilistic inference modes: (i) supervised classification using predefined ancestry labels, (ii) fraction-supervised modelling using continuous ancestry weights derived from reference populations, and (iii) fully unsupervised modelling in which latent population structure is learned directly from the genomic feature space without predefined ancestry labels.

### Construction of ancestry-informative k-mer resources

Ancestry-informative loci were derived from the GRCh38-integrated variant call (IVC) set generated through the reanalysis of the 1000 Genomes Project^19,20^. The source dataset consisted of the integrated phased biallelic SNV and INDEL call set generated from 2,548 unrelated individuals following realignment of the original 1000 Genomes sequencing data to the GRCh38 reference assembly.

Specifically, variants exhibiting population-specific allele frequencies exceeding 1% in at least one continental population were retained. This threshold was selected previously during ntRoot resource construction to enrich for population-informative variants while excluding extremely rare alleles that are less likely to contribute robust ancestry signal across sequencing datasets. The resulting filtered catalogue formed the ancestry-informative variant set used throughout KORUS model construction.

### Allele-centred k-mer generation

Unlike ntRoot, which identifies known population-informative variants in the query genome and performs ancestry inference using the allele frequencies associated with the identified variants in the IVC, KORUS uses k-mers derived from the IVC solely as quantitative signal generators. In brief, reference (REF) and alternate (ALT) allele k-mers were generated using a custom Perl utility that is included with the KORUS distribution (extract_allele_kmers.pl). For each variant, a fixed-length k-mer centred on the variant position was extracted from the GRCh38 reference sequence and corresponding REF and ALT k-mer representations were generated. The procedure supports single nucleotide variants (SNVs), insertions, deletions, and multi-allelic variants. For insertion and deletion alleles, flanking reference sequences were adjusted as necessary to ensure that all allele representations retained a constant k-mer length while remaining centred on the variant position. Two independent k-mer catalogues were generated using k = 45 and k = 55, each comprising ∼53.6M k-mers (Supplementary Table 1). All downstream analyses were performed independently for *k* = 45 and *k* = 55 to assess the effect of parameter *k* on KORUS ancestry inference.

### Reference panel construction

The KORUS reference panel used in this study consists of 643 genomes derived from the Simons Genome Diversity Project (SGDP, n = 279) and the 1000 Genomes Project (1kGP, n = 364)^1,2^. Combining SGDP and 1kGP increased both geographic diversity and representation of globally distributed populations while preserving compatibility with widely used population descriptors. Reference samples were not included in the blind-test cohort used for benchmarking (described below under **Benchmarking design**). Each reference genome was assigned a global ancestry label provided by the originating consortium, and the resulting ancestry composition is summarized in Supplementary Table 2. For hard-supervised decoding, all seven broad ancestry groups were represented: African (AFR), Admixed American (AMR), Central Asian (CAS), East Asian (EAS), European (EUR), Oceanian (OCE), and South Asian (SAS). For soft-supervised decoding, continuous ancestry fractions were obtained from prior ntRoot analyses^11^ of the same 643 reference genomes and used as ancestry weights during probabilistic decoding. Because these reference fractions were available only for the five continental super-populations represented by ntRoot, soft-supervised analyses were limited to African (AFR), Admixed American (AMR), East Asian (EAS), European (EUR), and South Asian (SAS).

### Bloom filter construction

Whole-genome sequencing reads were converted into probabilistic k-mer membership structures (i.e., Bloom filters) using the ntCard/ntStat ecosystem. Bloom filters represent k-mer presence using a probabilistic set-membership structure that permits false-positives but not false-negatives in membership queries. They provide a compact representation of sequencing-derived k-mer content, enabling rapid interrogation of large numbers of ancestry-informative k-mers while maintaining fixed memory footprints. For each sample, ntCard^34^ estimated the k-mer abundance spectrum and ntStat^35^ constructed a Bloom filter using the ntCard histogram. Bloom filters were generated using ntStat default parameters, corresponding to an expected false positive rate of approximately 0.001. Although the nominal false-positive rate was 0.001, realized false-positive rates may be lower depending on Bloom filter occupancy and sequencing-derived k-mer complexity. Ancestry-informative allele k-mers were queried against the sample Bloom filters (described above under **Allele-centred k-mer generation)**. Bloom filter generation and k-mer interrogation were performed using korusEncode, a Snakemake-based workflow^36^ distributed with KORUS. Briefly, korusEncode executes ntCard to estimate the k-mer abundance spectrum, constructs sample-specific Bloom filters using ntStat, and subsequently interrogates those Bloom filters using the ancestry-informative REF and ALT k-mer catalogues described above to yield a signal matrix.

### Variant-level ancestry signal extraction

For each ancestry-informative locus, KORUS queries the reference allele k-mer and all alternate allele k-mers defined in the ancestry-informative variant catalogue against the sample Bloom filter. Positive Bloom filter hits were accumulated separately for the reference allele representation and across all alternate allele representations, yielding *N_REF_* and *N_ALT_*, respectively. Because a locus may contain multiple alternate alleles, support was accumulated across all matching allele-specific k-mers rather than reduced to a binary presence/absence representation. To counter the influence of repetitive sequence content, Bloom filter false positives, and multi-allelic redundancy, evidence counts were variance-stabilized using logarithmic compression:

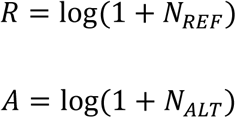

with *N_REF_* + *N_ALT_* ≥ 1. A normalized directional ancestry signal is then calculated as:

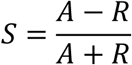

where S ranges from −1 to +1.

Positive values indicate stronger support for alternate-allele sequence content, whereas negative values indicate stronger support for reference-allele sequence content. Values near zero indicate balanced support for both allele representations. This formulation preserves the direction of allele support while normalizing for total signal intensity, thereby reducing sensitivity to local differences in k-mer density and sequencing depth.

### Genomic window aggregation

Variant-level signals were aggregated into non-overlapping genomic windows of 500 kb. Aggregation over fixed genomic windows reduces locus-level stochastic noise arising from sequencing coverage variation, Bloom filter uncertainty, and individual variant effects, while preserving regional ancestry structure. For each genomic window *w*, the window signal was computed as the arithmetic mean of all variant signals contained within that interval:

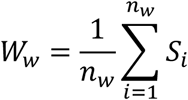

where *W_w_* denotes the aggregated ancestry signal for genomic window *w*, and *n_w_* denotes the number of informative loci within that window. When neither REF nor ALT evidence was detected at an informative locus, the locus signal was set to zero to avoid division by zero; this neutral value was included in the arithmetic mean used to calculate the window signal. Across the GRCh38 reference genome, this procedure yielded 5,465 non-overlapping windows. The resulting representation consists of a continuous genome-wide ancestry signal profile rather than a conventional genotype or variant matrix.

### Feature matrix construction

The aggregated window signals from all reference genomes are merged into a reference feature matrix in which columns and rows correspond to genomes and genomic windows, respectively. During decoding, query samples undergo the same transformation and are projected into the same feature space. Reference and query matrices are aligned by genomic window coordinates using identifiers of the form:

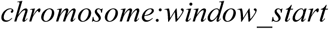

Only windows present in both reference and query datasets were retained for analysis.

### PCA-based local ancestry classification

For each focal genomic window, ancestry inference is performed using neighbouring genomic windows. Features within each neighbourhood are standardized and projected into principal component (PCA) space prior to probabilistic classification. PCA produces orthogonal latent variables that capture the dominant axes of variation within the local genomic feature space while removing linear correlation among genomic features. Mahalanobis distance estimation is therefore performed in a decorrelated representation rather than the original window space^37^. KORUS performs ancestry classification independently for each genomic window while incorporating a local neighbourhood of adjacent windows. For a target window *i*, neighbouring windows within a user-defined radius *r* were included, resulting in a local feature block spanning up to (*2r + 1*) adjacent windows. Unless otherwise specified, analyses were performed using the default radius of three windows. Incorporating neighboring windows increases local genomic context and reduces sensitivity to stochastic variation arising from individual windows. Reference and query matrices were standardized using z-score normalization and projected into PCA space. For each local ancestry model, the number of principal components (*n_pc_*) was selected dynamically as:

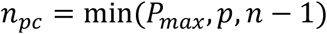

where *P_max_* is the user-specified maximum number of principal components (default = 10), *p* denotes the number of genomic features within the local window, and *n* denotes the number of reference samples used to construct the local model. This constraint ensures that the number of retained principal components does not exceed the dimensionality of the local feature space or the maximum rank supported by the centered reference matrix. The user-defined cap (*P_max_*) additionally limits the dimensionality of the representation used for Gaussian classification, helping avoid unnecessarily high-dimensional models when the available reference sample size is limited.

### Hard-supervised ancestry modelling

In hard-supervised mode, each reference genome contributes exclusively to a single ancestry class. For each ancestry class *c*, an ancestry-specific mean vector (μ_c_) and covariance matrix (Σ_c_) were estimated from the reference samples belonging to that class. Covariance matrices are regularized by adding:

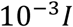

to enhance numerical stability and guarantee invertibility. For a query window represented by vector *z*, the squared Mahalanobis distance to ancestry class *c* is calculated as:

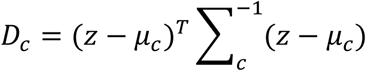

Squared Mahalanobis distances were converted into class-specific distance-derived scores, defined as:

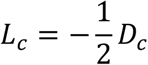

These scores were normalized across ancestry classes using a softmax transformation to obtain relative ancestry probabilities. Because this formulation omits the covariance log-determinant term, the scores are not full Gaussian log-likelihoods; rather, they quantify relative ancestry affinity based on covariance-adjusted distance.

### Soft-supervised ancestry modelling

In soft-supervised mode, reference genomes contribute fractional ancestry memberships supplied as ancestry-fraction weights. In this study, these weights were derived from prior ntRoot ancestry analyses of the reference panel. For ancestry *c*, weighted Gaussian parameters are estimated using ancestry fraction weights *w_ic_*. Weighted ancestry means are computed as:

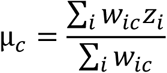

Weighted covariance matrices are estimated using weighted outer products and regularized identically to the hard-supervised model:

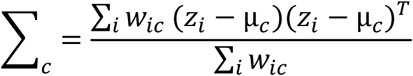

This formulation allows admixed reference genomes to contribute proportionally to multiple ancestry distributions, reducing dependence on hard ancestry labels and permitting reference genomes with admixed ancestry to contribute proportional information to multiple ancestry distributions.

### Unsupervised ancestry modelling

In unsupervised mode, no ancestry labels or ancestry fractions were provided. Reference genomes were standardized and projected into an orthogonal PCA-reduced feature space and partitioned into *K* clusters using K-means clustering. By default, clustering was performed using the first ten principal components, although this value could be modified by the user. The resulting cluster assignments were retained as pseudo-labels, and Gaussian ancestry models were subsequently estimated using the same procedure as the hard-supervised model. Posterior probabilities therefore represent affinity to inferred ancestry clusters rather than predefined ancestry categories. Unless otherwise specified, unsupervised analyses were performed using *K* = 5 clusters. The PCA used for unsupervised clustering serves only to generate the K-means cluster assignments used to initialize the ancestry models. It is distinct from the local PCA transformations used for window-level ancestry inference and the PCA visualizations generated for output reports.

### Probability estimation

Gaussian log-likelihood scores were converted into relative ancestry probabilities using a softmax transformation. The resulting posterior probabilities represent ancestry affinities for each genomic window. Window ancestry assignments are defined by the ancestry with maximal posterior probability:

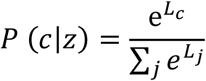

where *L_c_* denotes the Gaussian log-likelihood for ancestry class *c*, and the denominator sums over all ancestry classes *j*.

### Hidden Markov model smoothing

To reduce high-frequency ancestry switching, KORUS optionally applies a hidden Markov model to ancestry posterior probabilities. HMM smoothing was applied independently to each sample after posterior probability estimation. The model assumes that neighboring genomic windows are more likely to share the same ancestry state than distant windows and employs a transition matrix parameterized by a user-defined stay probability:

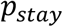

Transition probabilities are then defined as:

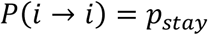

and

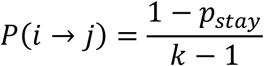

for all *i* ≠ *j*, assigning equal probabilities for transitioning to any other ancestry.

The most probable ancestry trajectory was recovered using the Viterbi algorithm. Unless otherwise specified, analyses used *p_stay_*=0.98, an empirically selected value that reduced high-frequency switching while preserving extended ancestry transitions. Softmax-normalized probabilities derived from the distance-based ancestry scores were used as HMM emission weights, allowing window-level ancestry affinities and neighborhood continuity to contribute to the final ancestry trajectory.

### Genome-wide ancestry fraction estimation

Final ancestry fractions were computed in two complementary ways. Hard ancestry fractions were calculated as the proportion of genomic windows assigned to each ancestry class following optional HMM smoothing. Posterior ancestry fractions were computed by averaging ancestry probability vectors across all genomic windows. Both outputs were reported for each sample.

### Benchmarking design

Independent validation was performed using 22 high-coverage whole-genome sequencing samples from the 1000 Genomes Project that were excluded from reference model construction (Supplementary Table 3). The benchmark cohort comprised individuals representing the five major 1000 Genomes Project superpopulations, including African (AFR, n = 4), Admixed American (AMR, n = 6), East Asian (EAS, n = 4), European (EUR, n = 4), and South Asian (SAS, n = 4). Raw 30× whole-genome sequencing data aligned to GRCh38 were obtained from the 1000 Genomes Project high-coverage data release. Individuals were selected at random within each superpopulation while ensuring no overlap with the KORUS reference panel. Consortium-provided superpopulation assignments were treated as the reference labels for benchmarking purposes.

Ancestry predictions generated by KORUS were compared against ntRoot^11^, SNVstory^18^, and ADMIXTURE^7^. Classification accuracy was calculated as the proportion of samples whose highest-probability ancestry assignment matched the corresponding 1000 Genomes Project superpopulation label. We also computed a normalized multiclass Brier score for each of the tools with the exception of SNVstory, which does not output estimated ancestral fractions. The Brier score falls within [0, 1], with values closer to 0 indicating higher concordance with ground truth labels. Concordance between ancestry fraction estimates was evaluated using coefficients of determination (R²) and concordance correlation coefficient (CCC).

### ntRoot ancestry inference

Global ancestry inference was performed using ntRoot v1.1.6 as previously described^11^. Sequencing reads from each benchmark sample were analyzed directly using the ntRoot alignment-free workflow with default ancestry resources derived from the 1000 Genomes Project. Analyses were performed independently using k-mer lengths of 45 and 55. For each sample, the reported ancestry fractions and highest-probability ancestry assignment were retained for downstream comparisons with KORUS.

### SNVstory ancestry inference

SNVstory requires variant-based input and therefore sequencing reads were first aligned to the GRCh38 human reference genome using BWA-MEM^38^. Variant discovery was subsequently performed using the Genome Analysis Toolkit (GATK) Best Practices workflow^39^. Briefly, variants were called independently for each chromosome using GATK HaplotypeCaller and merged into a genome-wide variant call set using BCFtools^40^. The resulting variant call files were analyzed using SNVstory v3.0.2 in whole-genome sequencing mode with the GRCh38 reference configuration. Global ancestry predictions were extracted from the continental ancestry probabilities reported by SNVstory, and the ancestry with the highest posterior probability was used as the primary classification.

### ADMIXTURE ancestry inference

ADMIXTURE analyses were performed using a supervised reference-panel design. Reference genotype data were obtained from the high-coverage 1000 Genomes Project call set after excluding all benchmark samples. Variant calls generated for each benchmark individual were merged with the reference genotype panel using BCFtools and filtered to retain autosomal biallelic variants. The merged genotype dataset was converted to PLINK format and analyzed using ADMIXTURE v1.3.0 in supervised mode^41^. Ancestry fractions reported by ADMIXTURE were used for downstream comparisons with KORUS and ntRoot. For classification accuracy calculations, the ancestry component with the largest estimated fraction was treated as the predicted ancestry assignment.

### Ancient genome analyses

To evaluate model behaviour beyond the diversity represented in contemporary human reference panels, KORUS was applied to three ancient genomes: the Denisovan genome^42^ (ERP001519), the Altai Neanderthal genome^43^ (ERP002097), and the high-coverage Tyrolean Iceman genome^29^ (ERP141520). These analyses served as qualitative demonstrations of ancestry-projection behaviour under substantial divergence from the modern reference panel, rather than as formal classification benchmarks.

### Computational resource utilization

Resource utilization was measured using the UNIX time utility. Wall-clock runtime and peak resident memory consumption were recorded for all major processing stages, including ntCard, ntStat, korusEncode, korusMatrix.py, korusDecode.py, ntRoot, SNVstory, and ADMIXTURE. KORUS analyses were performed on a server equipped with 144 Intel Xeon Gold 6150 processors and 2.1 TB RAM using 48 computational threads. Comparative benchmarking of ntRoot, SNVstory, and ADMIXTURE was performed on a server equipped with 144 Intel Xeon Gold 6254 processors and 2.9 TB RAM.

### Software implementation and availability

KORUS is implemented in Python and runs on Linux and Mac OS. The framework comprises modular encoding and decoding components (korusEncode, korusMatrix.py, and korusDecode.py) together with an end-to-end execution pipeline named korus. Interactive visualizations of PCA-derived population structure and local ancestry trajectories were generated using Plotly. Source code, documentation, and example datasets are freely available under the GPLv3 license at https://github.com/BirolLab/korus.

## Author contributions

Conceptualization: RLW, IB. Data curation: RLW. Formal analysis: RLW, LC. Funding acquisition: RLW, IB. Investigation: RLW. Methodology: RLW. Project administration: RLW, IB. Resources: RLW. Software: RLW. Supervision: RLW, IB. Validation: RLW. Visualization: RLW, LC, Writing – original draft: RLW, LC, Writing – review & editing: RLW, LC, IB.

## Data availability

The data supporting this study are publicly available from the repositories and sources described below. The KORUS reference panel comprises 643 whole-genome sequences from the 1000 Genomes Project (1kGP; n = 364) and the Simons Genome Diversity Project (SGDP; n = 279). The 1kGP reference genomes were obtained from the International Genome Sample Resource (IGSR) 30× GRCh38 dataset (https://www.internationalgenome.org/data-portal/data-collection/30x-grch38), with sample/accession identifiers provided in Supplementary Table 13. The SGDP genome sequencing data are available through the European Nucleotide Archive (ENA) under study accession PRJEB9586 (https://www.ebi.ac.uk/ena/browser/view/PRJEB9586). The GRCh38 reference assembly (GCF_000001405.15) is available from NCBI (https://www.ncbi.nlm.nih.gov/datasets/genome/GCF_000001405.15/). The 1kGP variant representations (i.e., VCFs) used to derive ancestry-informative k-mers are available through IGSR (https://www.internationalgenome.org/data-portal/data-collection/30x-grch38). Both files are also archived on Zenodo (https://doi.org/10.5281/zenodo.10869033). The independent benchmark cohort comprises 22 whole-genome sequencing samples from 1kGP that were excluded from reference model construction. Their ENA accession numbers and corresponding sample identifiers are provided in Supplementary Table 3. The ancient genome sequencing data used for ancestry projection are available under the following ENA (https://www.ebi.ac.uk/ena/) study accessions: Denisovan (ERP001519), Altai Neanderthal (ERP002097), and Tyrolean Iceman (ERP141520).

## References

1. Auton, A. et al. A global reference for human genetic variation. Nature 526, 68–74 (2015).

2. Mallick, S. et al. The Simons Genome Diversity Project: 300 genomes from 142 diverse populations. Nature 538, 201–206 (2016).

3. Conley, A. B. et al. Rye: genetic ancestry inference at biobank scale. Nucleic Acids Res. 51, e44 (2023).

4. Gouveia, M. H. et al. Subcontinental genetic variation in the All of Us Research Program: Implications for biomedical research. Am. J. Hum. Genet. 112, 1286–1301 (2025).

5. Warren, R. L., Coombe, L. & Birol, I. Concordance and divergence between self-declared ancestry and genome-derived ancestry composition in 10 250 participants from the HostSeq cohort. Genome 69, 1–12 (2026).

6. Pritchard, J. K., Stephens, M. & Donnelly, P. Inference of population structure using multilocus genotype data. Genetics 155, 945–959 (2000).

7. Alexander, D. H., Novembre, J. & Lange, K. Fast model-based estimation of ancestry in unrelated individuals. Genome Res. 19, 1655–1664 (2009).

8. Price, A. L. et al. Principal components analysis corrects for stratification in genome-wide association studies. Nat. Genet. 38, 904–909 (2006).

9. Maples, B. K., Gravel, S., Kenny, E. E. & Bustamante, C. D. RFMix: a discriminative modeling approach for rapid and robust local-ancestry inference. Am. J. Hum. Genet. 93, 278–288 (2013).

10. Browning, S. R., Waples, R. K. & Browning, B. L. Fast, accurate local ancestry inference with FLARE. Am. J. Hum. Genet. 110, 326–335 (2023).

11. Warren, R. L., Coombe, L., Wong, J., Kazemi, P. & Birol, I. ntRoot: computational inference of human ancestry at scale from genomic data. Bioinforma. Adv. 5, vbaf287 (2025).

12. Gibbs, R. A. et al. The International HapMap Project. Nature 426, 789–796 (2003).

13. Committee on the Use of Race, Ethnicity, and Ancestry as Population Descriptors in Genomics Research et al. Using Population Descriptors in Genetics and Genomics Research: A New Framework for an Evolving Field. 26902 (National Academies Press, Washington, D.C., 2023). doi:10.17226/26902.

14. Bloom, B. H. Space/time trade-offs in hash coding with allowable errors. Commun. ACM 13, 422–426 (1970).

15. Warren, R. L. et al. LINKS: Scalable, alignment-free scaffolding of draft genomes with long reads. Gigascience 4, s13742-015-0076–3 (2015).

16. Jackman, S. D. et al. ABySS 2.0: resource-efficient assembly of large genomes using a Bloom filter. Genome Res. 27, 768–777 (2017).

17. Rahman, A., Hallgrímsdóttir, I., Eisen, M. & Pachter, L. Association mapping from sequencing reads using k-mers. eLife 7, e32920 (2018).

18. Bollas, A. E. et al. SNVstory: inferring genetic ancestry from genome sequencing data. BMC Bioinformatics 25, 76 (2024).

19. Zheng-Bradley, X. et al. Alignment of 1000 Genomes Project reads to reference assembly GRCh38. GigaScience 6, (2017).

20. Lowy-Gallego, E. et al. Variant calling on the GRCh38 assembly with the data from phase three of the 1000 Genomes Project. Wellcome Open Res. 4, 50 (2019).

21. Zimin, A. V. et al. A reference-quality, fully annotated genome from a Puerto Rican individual. Genetics 220, iyab227 (2022).

22. Gourraud, P.-A. et al. HLA diversity in the 1000 genomes dataset. PloS One 9, e97282 (2014).

23. Fedorova, L., Qiu, S., Dutta, R. & Fedorov, A. Atlas of Cryptic Genetic Relatedness Among 1000 Human Genomes. Genome Biol. Evol. 8, 777–790 (2016).

24. Shi, G. & Kuang, Q. Ancestral spectrum analysis with population-specific variants. Front. Genet. 12, 724638 (2021).

25. Gravel, S. et al. Reconstructing Native American migrations from whole-genome and whole-exome data. PLoS Genet. 9, e1004023 (2013).

26. Campbell, M. C. & Tishkoff, S. A. African Genetic Diversity: Implications for Human Demographic History, Modern Human Origins, and Complex Disease Mapping. Annu. Rev. Genomics Hum. Genet. 9, 403–433 (2008).

27. Keller, A. et al. New insights into the Tyrolean Iceman’s origin and phenotype as inferred by whole-genome sequencing. Nat. Commun. 3, 698 (2012).

28. Croze, M. et al. Genomic diversity and structure of prehistoric alpine individuals from the Tyrolean Iceman’s territory. Nat. Commun. 16, 6431 (2025).

29. Wang, K. et al. High-coverage genome of the Tyrolean Iceman reveals unusually high Anatolian farmer ancestry. Cell Genomics 3, 100377 (2023).

30. Patterson, N., Price, A. L. & Reich, D. Population structure and eigenanalysis. PLoS Genet. 2, e190 (2006).

31. Lawson, D. J., van Dorp, L. & Falush, D. A tutorial on how not to over-interpret STRUCTURE and ADMIXTURE bar plots. Nat. Commun. 9, 3258 (2018).

32. Li, J. Z. et al. Worldwide human relationships inferred from genome-wide patterns of variation. Science 319, 1100–1104 (2008).

33. Novembre, J. et al. Genes mirror geography within Europe. Nature 456, 98–101 (2008).

34. Mohamadi, H., Khan, H. & Birol, I. ntCard: a streaming algorithm for cardinality estimation in genomics data. Bioinformatics 33, 1324–1330 (2017).

35. Kazemi, P., Coombe, L., Warren, R. L. & Birol, I. ntStat: k-mer characterization using occurrence statistics in raw sequencing data. PLOS Comput. Biol. 22, e1014158 (2026).

36. Mölder, F. et al. Sustainable data analysis with Snakemake. F1000Research 10, 33 (2021).

37. Mahalanobis, P. C. Reprint of: Mahalanobis, P.C. (1936) ‘On the Generalised Distance in Statistics.’ Sankhya A 80, 1–7 (2018).

38. Li, H. Aligning sequence reads, clone sequences and assembly contigs with BWA-MEM. (2013) doi:10.48550/ARXIV.1303.3997.

39. Van der Auwera, G. A. & O’Connor, B. D. Genomics in the Cloud: Using Docker, GATK, and WDL in Terra. (O’Reilly Media, 2020).

40. Danecek, P. et al. Twelve years of SAMtools and BCFtools. GigaScience 10, (2021).

41. Chang, C. C. et al. Second-generation PLINK: rising to the challenge of larger and richer datasets. GigaScience 4, s13742-015-0047–8 (2015).

42. Meyer, M. et al. A High-Coverage Genome Sequence from an Archaic Denisovan Individual. Science 338, 222–226 (2012).

43. Prüfer, K. et al. The complete genome sequence of a Neanderthal from the Altai Mountains. Nature 505, 43–49 (2014).

