## Supplementary Material for "KORUS: probabilistic ancestry inference beyond variant calls"

### Supplementary Tables

**Supplementary Table 1. Summary of the allele-centered k-mer reference catalogues used by KORUS.** The numbers of reference (REF) and alternate (ALT) allele k-mers generated for k = 45 and k = 55 are shown. These catalogues were used independently throughout all downstream analyses to evaluate the effect of k-mer length on ancestry inference.

| k-mer size | REF k-mers | ALT k-mers |
|---|---|---|
| **45** | 26,749,020 | 26,823,406 |
| **55** | 26,755,870 | 26,830,256 |

**Supplementary Table 2. Global ancestry composition of the KORUS reference panel.** The combined reference panel includes 643 genomes from the Simons Genome Diversity Project (SGDP) and the 1000 Genomes Project (1kGP), stratified according to the ancestry labels provided by the originating consortia. These labels served as the reference classes for hard-supervised ancestry inference.

| Ancestry hard label (provided by SGDP/1kGP) | Sample count |
|---|---|
| **African (AFR)** | 137 |
| **Admixed American (AMR)** | 82 |
| **Central Asian (CAS)** | 27 |
| **East Asian (EAS)** | 118 |
| **European (EUR)** | 145 |
| **Oceanian (OCE)** | 25 |
| **South Asian (SAS)** | 109 |
| **Total** | 643 |

**Supplementary Table 3. Composition of the independent benchmark cohort used for performance evaluation.** The validation set comprises 22 high-coverage whole-genome sequencing samples from the 1000 Genomes Project (1kGP) that were excluded from reference model construction. Samples represent the five major 1kGP superpopulations (African (AFR), Admixed American (AMR), East Asian (EAS), European (EUR), and South Asian (SAS)) and were used as the blind-test cohort for benchmarking KORUS, ntRoot, SNVstory, and ADMIXTURE. Letter codes correspond to sample identifiers used in figures throughout the manuscript.

| Accession | Sample | Letter code | Superpopulation code* |
|---|---|---|---|
| **ERR3239665** | NA18865 | a | AFR |
| **ERR3239767** | NA19449 | b | AFR |
| **ERR3239986** | NA20314 | c | AFR^ (AMR) |
| **ERR3241775** | NA20321 | d | AFR |
| **ERR3241809** | HG01069 | e | AMR |
| **ERR3241810** | HG01070 | f | AMR |
| **ERR3241845** | HG01174 | g | AMR |
| **ERR3241865** | HG01254 | h | AMR |
| **ERR3242010** | HG01947 | i | AMR |
| **ERR3988858** | HG01243 | j | AMR^ (AFR) |
| **ERR3239524** | NA18627 | k | EAS |
| **ERR3240184** | HG00443 | l | EAS |
| **ERR3241741** | HG00684 | m | EAS |
| **ERR3242062** | HG01595 | n | EAS |
| **ERR3240246** | HG00338 | o | EUR |
| **ERR3240253** | HG00349 | p | EUR |
| **ERR3240254** | HG00350 | q | EUR |
| **ERR3240258** | HG00356 | r | EUR |
| **ERR3242841** | HG03808 | s | SAS |
| **ERR3242871** | HG03672 | t | SAS |
| **ERR3243086** | HG04189 | u | SAS |
| **ERR3243087** | HG04194 | v | SAS |

*From 1kGP. ^Identified in studies as mislabeled, with the likely correct GAI in parentheses [1–4].

**Supplementary Table 4. Ancestry fractions from running KORUS in hard-supervised mode on 22 benchmarking samples.**
See excel spreadsheet korusMS_supplementaryTablesS4-S8andS13.xlsx

**Supplementary Table 5. Ancestry fractions from running KORUS in soft-supervised mode on 22 benchmarking samples.**
See excel spreadsheet korusMS_supplementaryTablesS4-S8andS13.xlsx

**Supplementary Table 6. Ancestry fractions from running KORUS in unsupervised mode on 22 benchmarking samples.** The clusters were assigned to ancestries based on the majority label of the reference samples making up that cluster.
See excel spreadsheet korusMS_supplementaryTablesS4-S8andS13.xlsx

**Supplementary Table 7. Ancestry fractions from running ntRoot on 22 benchmarking samples.**
See excel spreadsheet korusMS_supplementaryTablesS4-S8andS13.xlsx

**Supplementary Table 8. Ancestry fractions from running ADMIXTURE on 22 benchmarking samples.**
See excel spreadsheet korusMS_supplementaryTablesS4-S8andS13.xlsx

**Supplementary Table 9. Accuracy and normalized multiclass Brier scores from ancestry predictions using KORUS, ntRoot, ADMIXTURE and SNVstory.** Global ancestry inference was performed using whole genome sequencing reads from 22 human samples. For KORUS (k=55), ntRoot and ADMIXTURE, the global ancestry label was assigned to the super-population with the largest ancestry fraction, while for SNVstory, the label with the highest probability was chosen. The classification was considered correct if it matched the 1kGP label, and the accuracy was calculated as the percentage of correctly classified samples.

| Tool | Number of correctly classified samples | Accuracy (%) | Brier Score |
|---|---|---|---|
| **KORUS Hard** | 16 | 72.70 | 0.168 |
| **KORUS Soft** | 18 | 81.80 | 0.137 |
| **KORUS Unsupervised** | 15 | 68.20 | 0.236 |
| **ntRoot** | 20 | 90.90 | 0.131 |
| **ADMIXTURE** | 20 | 90.90 | 0.097 |
| **SNVstory** | 22 | 100.00 | NA |

**Supplementary Table 10. Global ancestry inference results from running KORUS, ntRoot, ADMIXTURE and SNVstory on 22 samples.** Ground truth super-populations were determined from 1kGP. The bolded cells indicate an incorrect classification (different from the ground truth). The two samples indicated with an asterisk (NA20314 and HG01243) have been identified in previous studies as being potentially mislabeled, with data indicating that their more appropriate super-population may be AMR and AFR, respectively.

| Accession | Sample ID | Ground truth label | KORUS Hard | KORUS Soft | KORUS Unsupervised | ntRoot | ADMIXTURE | SNVstory |
|---|---|---|---|---|---|---|---|---|
| ERR3239665 | NA18865 | AFR | AFR | AFR | AFR | AFR | AFR | AFR |
| ERR3239767 | NA19449 | AFR | AFR | AFR | AFR | AFR | AFR | AFR |
| ERR3239986 | NA20314* | AFR | **AMR** | **AMR** | **EUR** | **AMR** | **AMR** | AFR |
| ERR3241775 | NA20321 | AFR | AFR | AFR | AFR | AFR | AFR | AFR |
| ERR3241809 | HG01069 | AMR | **EUR** | AMR | **EUR** | AMR | AMR | AMR |
| ERR3241810 | HG01070 | AMR | **EUR** | **EUR** | **EUR** | AMR | AMR | AMR |
| ERR3241845 | HG01174 | AMR | **EUR** | **EUR** | **EUR** | AMR | AMR | AMR |
| ERR3241865 | HG01254 | AMR | **EUR** | AMR | **EUR** | AMR | AMR | AMR |
| ERR3242010 | HG01947 | AMR | AMR | AMR | **EUR** | AMR | AMR | AMR |
| ERR3988858 | HG01243* | AMR | **AFR** | **AFR** | **AFR** | **AFR** | **AFR** | AMR |
| ERR3239524 | NA18627 | EAS | EAS | EAS | EAS | EAS | EAS | EAS |
| ERR3240184 | HG00443 | EAS | EAS | EAS | EAS | EAS | EAS | EAS |
| ERR3241741 | HG00684 | EAS | EAS | EAS | EAS | EAS | EAS | EAS |
| ERR3242062 | HG01595 | EAS | EAS | EAS | EAS | EAS | EAS | EAS |
| ERR3240246 | HG00338 | EUR | EUR | EUR | EUR | EUR | EUR | EUR |
| ERR3240253 | HG00349 | EUR | EUR | EUR | EUR | EUR | EUR | EUR |
| ERR3240254 | HG00350 | EUR | EUR | EUR | EUR | EUR | EUR | EUR |
| ERR3240258 | HG00356 | EUR | EUR | EUR | EUR | EUR | EUR | EUR |
| ERR3242841 | HG03808 | SAS | SAS | SAS | SAS | SAS | SAS | SAS |
| ERR3242871 | HG03672 | SAS | SAS | SAS | SAS | SAS | SAS | SAS |
| ERR3243086 | HG04189 | SAS | SAS | SAS | SAS | SAS | SAS | SAS |
| ERR3243087 | HG04194 | SAS | SAS | SAS | SAS | SAS | SAS | SAS |

**Supplementary Table 11. Accuracy and normalized multiclass Brier scores from ancestry prediction using KORUS, ntRoot, ADMIXTURE and SNVstory with adjusted ground truth labels.** Global ancestry inference was performed using whole genome sequencing reads from 22 human samples. For KORUS (k=55), ntRoot and ADMIXTURE, the global ancestry label was assigned to the super-population with the largest ancestry fraction, while for SNVstory, the label with the highest probability was chosen. The ground truth labels were taken from 1kGP, with the exception of samples HG01243 (relabeled to AFR) and NA20314 (relabeled to AMR), which past studies have suggested were mislabeled [1–4].

| Tool | Number of correctly classified samples | Accuracy (%) | Brier score |
|---|---|---|---|
| **KORUS Hard** | 18 | 81.80 | 0.115 |
| **KORUS Soft** | 20 | 90.90 | 0.069 |
| **KORUS Unsupervised** | 16 | 72.70 | 0.198 |
| **ntRoot** | 22 | 100.00 | 0.083 |
| **ADMIXTURE** | 22 | 100.00 | 0.035 |
| **SNVstory** | 20 | 90.90 | NA |

**Supplementary Table 12. Correlations between ancestry fractions produced by KORUS, ntRoot and ADMIXTURE.** The $R^2$ and CCC (concordance correlation coefficient) statistics were computed across all 22 blind test samples.

| Comparison | Ancestry | $R^2$ | CCC |
|---|---|---|---|
| **KORUS Hard vs. Soft** | AFR | 0.990 | 0.989 |
| **KORUS Hard vs. Soft** | AMR | 0.908 | 0.861 |
| **KORUS Hard vs. Soft** | EAS | 0.996 | 0.977 |
| **KORUS Hard vs. Soft** | EUR | 0.958 | 0.923 |
| **KORUS Hard vs. Soft** | SAS | 0.989 | 0.988 |
| **KORUS Hard k55 vs. k45** | AFR | 1.000 | 1.000 |
| **KORUS Hard k55 vs. k45** | AMR | 0.997 | 0.998 |
| **KORUS Hard k55 vs. k45** | CAS | 0.845 | 0.903 |
| **KORUS Hard k55 vs. k45** | EAS | 0.999 | 1.000 |
| **KORUS Hard k55 vs. k45** | EUR | 0.998 | 0.999 |
| **KORUS Hard k55 vs. k45** | OCE | 0.708 | 0.749 |
| **KORUS Hard k55 vs. k45** | SAS | 0.998 | 0.999 |
| **KORUS Soft k55 vs. k45** | AFR | 1.000 | 1.000 |
| **KORUS Soft k55 vs. k45** | AMR | 0.998 | 0.998 |
| **KORUS Soft k55 vs. k45** | EAS | 1.000 | 1.000 |
| **KORUS Soft k55 vs. k45** | EUR | 0.998 | 0.999 |
| **KORUS Soft k55 vs. k45** | SAS | 0.999 | 0.999 |
| **KORUS Hard k55 vs. KORUS Unsupervised** | AFR | 1.000 | 1.000 |
| **KORUS Hard k55 vs. KORUS Unsupervised** | AMR | 0.928 | 0.554 |
| **KORUS Hard k55 vs. KORUS Unsupervised** | EAS | 0.996 | 0.987 |
| **KORUS Hard k55 vs. KORUS Unsupervised** | EUR | 0.925 | 0.903 |
| **KORUS Hard k55 vs. KORUS Unsupervised** | OCE | 0.274 | 0.357 |
| **KORUS Hard vs. ntRoot** | AFR | 0.969 | 0.969 |
| **KORUS Hard vs. ntRoot** | AMR | 0.665 | 0.767 |
| **KORUS Hard vs. ntRoot** | EAS | 0.976 | 0.912 |
| **KORUS Hard vs. ntRoot** | EUR | 0.916 | 0.800 |
| **KORUS Hard vs. ntRoot** | SAS | 0.973 | 0.983 |
| **KORUS Soft vs. ntRoot** | AFR | 0.981 | 0.986 |
| **KORUS Soft vs. ntRoot** | AMR | 0.825 | 0.878 |
| **KORUS Soft vs. ntRoot** | EAS | 0.979 | 0.970 |
| **KORUS Soft vs. ntRoot** | EUR | 0.979 | 0.951 |
| **KORUS Soft vs. ntRoot** | SAS | 0.983 | 0.990 |
| **KORUS Hard vs. ADMIXTURE** | AFR | 0.977 | 0.983 |
| **KORUS Hard vs. ADMIXTURE** | AMR | 0.844 | 0.809 |
| **KORUS Hard vs. ADMIXTURE** | EAS | 0.989 | 0.916 |
| **KORUS Hard vs. ADMIXTURE** | EUR | 0.854 | 0.884 |
| **KORUS Hard vs. ADMIXTURE** | SAS | 0.987 | 0.941 |
| **KORUS Soft vs. ADMIXTURE** | AFR | 0.988 | 0.993 |
| **KORUS Soft vs. ADMIXTURE** | AMR | 0.894 | 0.902 |
| **KORUS Soft vs. ADMIXTURE** | EAS | 0.993 | 0.977 |
| **KORUS Soft vs. ADMIXTURE** | EUR | 0.950 | 0.950 |
| **KORUS Soft vs. ADMIXTURE** | SAS | 0.988 | 0.944 |

| | | | |
|---|---|---|---|
| **ntRoot vs. ADMIXTURE** | AFR | 0.995 | 0.989 |
| **ntRoot vs. ADMIXTURE** | AMR | 0.895 | 0.887 |
| **ntRoot vs. ADMIXTURE** | EAS | 0.987 | 0.991 |
| **ntRoot vs. ADMIXTURE** | EUR | 0.981 | 0.886 |
| **ntRoot vs. ADMIXTURE** | SAS | 0.984 | 0.953 |
| **ntRoot k55 vs. k45** | AFR | 0.995 | 0.980 |
| **ntRoot k55 vs. k45** | AMR | 0.987 | 0.991 |
| **ntRoot k55 vs. k45** | EAS | 0.998 | 0.997 |
| **ntRoot k55 vs. k45** | EUR | 0.999 | 0.961 |
| **ntRoot k55 vs. k45** | SAS | 0.999 | 0.996 |

**Supplementary Table 13. Samples used to build the KORUS reference panel.**
See excel spreadsheet korusMS_supplementaryTablesS4-S8andS13.xlsx

### Supplementary Figures

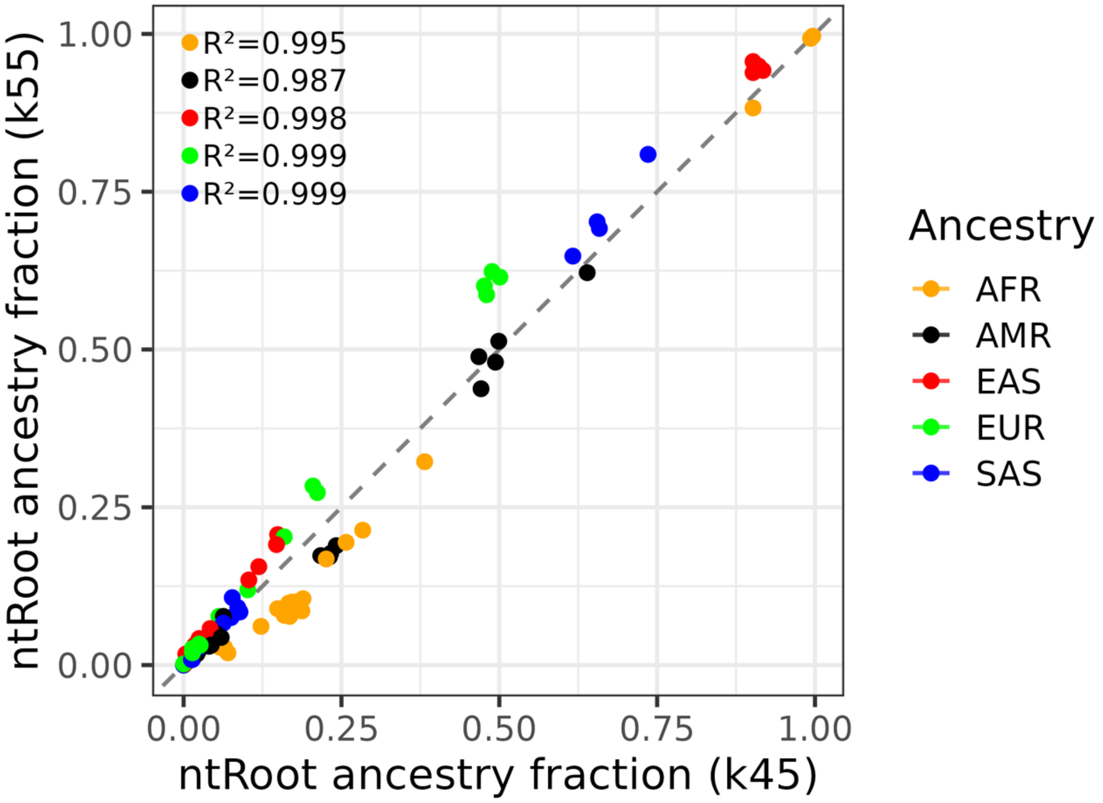

**Supplementary Figure 1. Concordance of ntRoot fractional ancestry predictions using k=45 and k=55.**

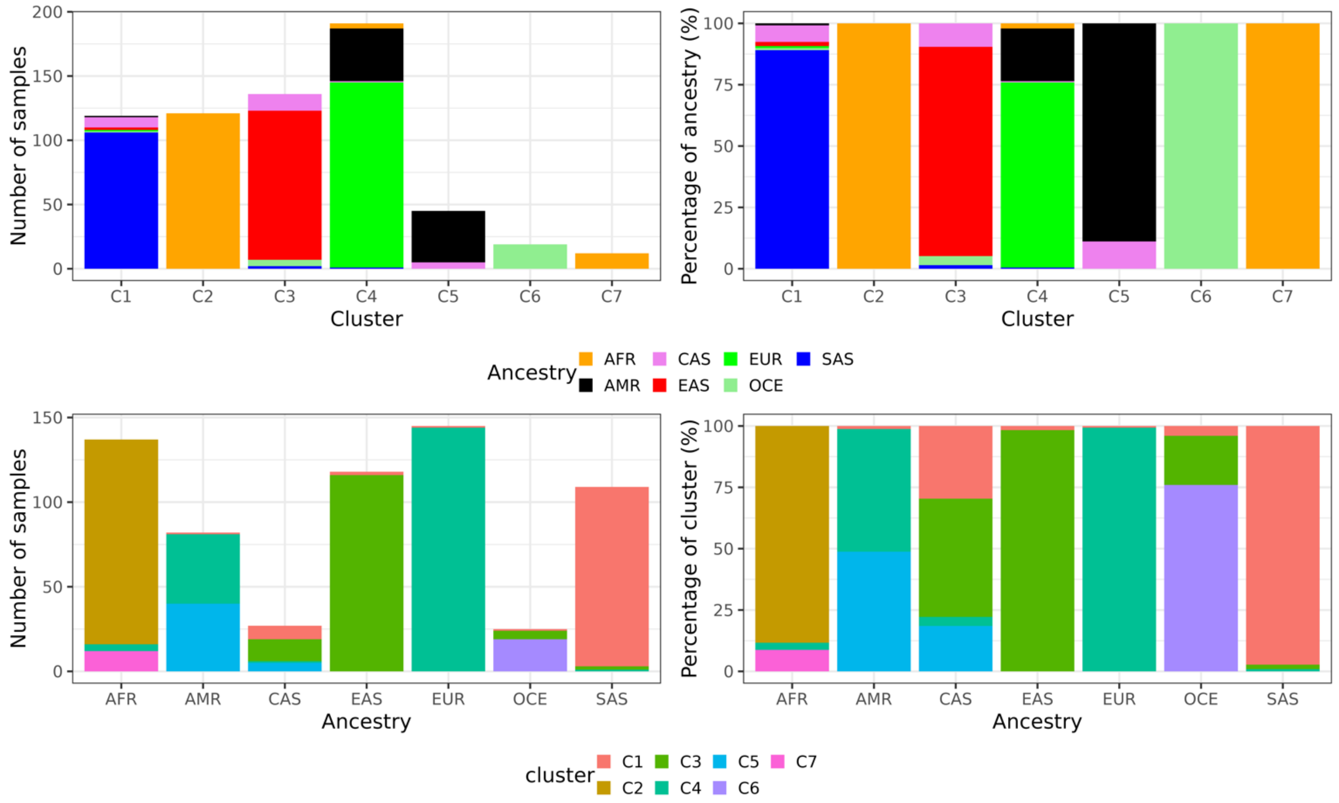

**Supplementary Figure 2. Unsupervised recovery of population structure by KORUS using k=45.** Characterization of unsupervised clustering of the reference panel using seven K-means clusters. Cluster composition is summarized for independent analyses using a k-mer length of 45. Stacked bar plots illustrate the distribution of known reference ancestries within each inferred cluster and the reciprocal distribution of ancestry groups across clusters.

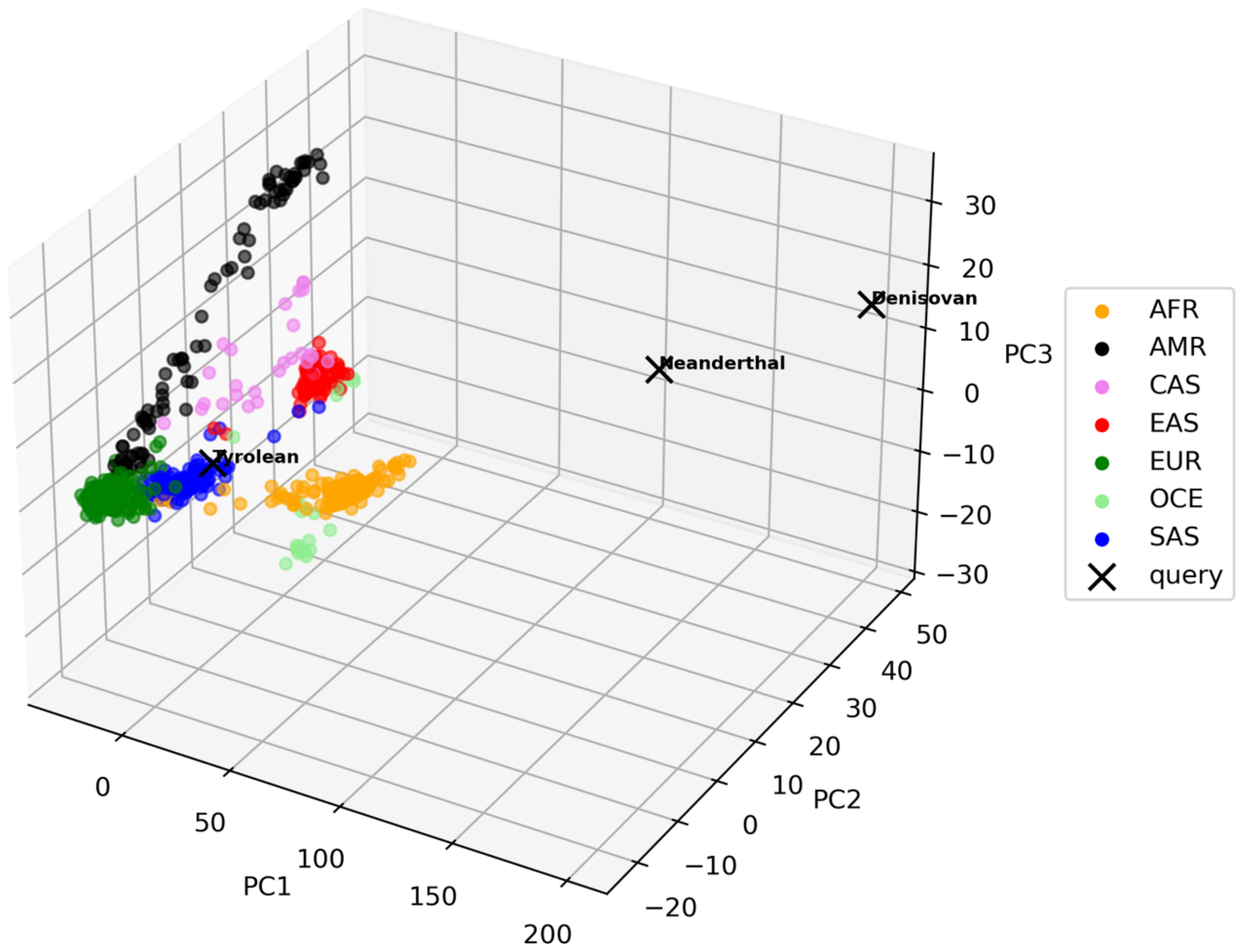

**Supplementary Figure 3. Projection of ancient human and archaic hominin genomes onto the KORUS hard-supervised ancestry reference space.** Three-dimensional principal component analysis (PCA) projection of the hard-supervised KORUS reference model ($k = 45$) showing the modern reference populations including African (AFR), Admixed American (AMR), Central Asian (CAS), East Asian (EAS), European (EUR), Oceanian (OCE), and South Asian (SAS) together with the Tyrolean Iceman (Ötzi), Neanderthal, and Denisovan genomes (marked with ×). Modern reference samples are coloured according to their assigned ancestry groups. The Tyrolean genome projects within the modern human genomic manifold with strongest affinity to European populations, whereas the Neanderthal and Denisovan genomes occupy positions well outside the distribution of present-day human variation. These projections illustrate that KORUS represents evolutionarily divergent genomes according to their proximity to the learned modern genomic manifold rather than assigning biologically meaningful continental ancestry to genomes that fall outside the distribution of the reference panel.

### References

1. Gourraud, P.-A. *et al.* HLA diversity in the 1000 genomes dataset. *PloS one* **9**, e97282 (2014).
2. Fedorova, L., Qiu, S., Dutta, R. & Fedorov, A. Atlas of Cryptic Genetic Relatedness Among 1000 Human Genomes. *Genome Biology and Evolution* **8**, 777–790 (2016).
3. Shi, G. & Kuang, Q. Ancestral spectrum analysis with population-specific variants. *Frontiers in Genetics* **12**, 724638 (2021).
4. Zimin, A. V. *et al.* A reference-quality, fully annotated genome from a Puerto Rican individual. *Genetics* **220**, iyab227 (2022).
